# Hybrid modelling and transfer learning for Bayesian optimisation of yeast protein production from food waste substrates

**DOI:** 10.64898/2026.07.24.740460

**Authors:** Alexander L. Bowler, Nasser Alkhulaifi, Sarah Bowler, Joanna H. Sier, Célia Ferreira, Darren Greetham, Jordan Pennells, Kai Knoerzer, Nicholas J. Watson

## Abstract

Food production is a significant contributor to global greenhouse gas emissions and deforestation, exacerbated by substantial food waste. Converting food waste into yeast protein offers a sustainable solution to enhance food security and contribute to a circular economy. However, due to the diverse and variable nature of food waste substrates, numerous experimental trials are required to optimise the preprocessing steps, yeast strain selection, nutrient addition, and fermentation conditions. This study presents a hybrid modelling approach where data-driven machine learning is used to predict microbial growth kinetics from process parameters. The hybrid model was trained on a comprehensive dataset consisting of 963 fermentation experiments from 55 publications, enabling transfer learning across 46 yeast strains and 79 food waste substrates. The hybrid modelling method was integrated with Bayesian optimisation, a sequential strategy to optimise expensive-to-evaluate functions, to efficiently maximise yeast biomass growth from different food waste substrates. The utility of the hybrid model was evaluated using five test datasets selected from previous literature and was shown to facilitate an average reduction of 66% in the number of experimental trials required to identify optimal fermentation conditions compared to without using the hybrid model. This proved that the transfer of knowledge between yeast strains and food wastes improved the optimisation efficiency of real, previously published datasets compared to traditional optimisation methods. The novelty and contributions of this study include the collation of the extensive dataset, provided as supplementary material; and the demonstration that transfer learning by training the hybrid model on this heterogeneous dataset can improve the optimisation efficiency for yeast biomass growth on new strains and substrates.

## 1 Introduction

Food production contributes approximately 30% of global Greenhouse Gas (GHG) emissions (Ritchie, 2021) with animal-sourced foods accounting for half of this (Xu et al., 2021). Furthermore, agricultural expansion drives around 90% of global deforestation, mainly for livestock grazing and feed production (FAO, 2021). Meanwhile, around one third of all food produced is wasted, equating to 8–10% of global GHG emissions and requiring 30% of agricultural land (UNEP, 2024). This inefficiency coexists with one third of people experiencing food insecurity (UNEP, 2024). With the world’s population projected to rise to 9.7 billion by 2050, food production must increase by up to 56% over 2010 levels (van Dijk et al., 2021). Demand for protein is also rising with estimates suggesting that demand for animal-derived protein could double by 2050 (Henchion et al., 2017). Therefore, alternative protein sources are needed to produce sufficient protein for a growing population, but to do so without worsening environmental problems.

Converting agri-food waste to microbial protein presents a promising solution to address these challenges of food security and environmental concerns. It has been estimated that utilisation of available agri-food waste streams could produce enough microbial protein to meet the global protein demand (Piercy et al., 2022). Moreover, valorisation of food side streams into microbial protein utilises nutrients that would otherwise have been wasted and contributes to the development of a circular economy. Yeast is a microorganism that has been grown to produce single cell protein in many previous studies by utilising nutrients from food waste streams. For example, *Saccharomyces cerevisiae* has previously been grown using waste vinasse (Dos Reis et al., 2019), fruit (Sawsan et al., 2021), leafy vegetables (Choi and Park, 2003), root vegetables (Tian et al., 2023), and whey (Trigueros et al., 2016); *Candida utilis* using waste molasses (Lee and Kim, 2001), vinasse (Tauk, 1982), fruit (Nigam, 1998), leafy vegetables (Choi and Park, 2003), and root vegetables (Ezekiel and Aworh, 2018); and *Yarrowia lipolytica* on waste bread hydrolysate (Carsanba et al., 2023). However, to maximise the economic and environmental sustainability of producing yeast protein from food waste streams, the preprocessing steps, yeast strain selection, nutrient addition, and fermentation conditions must be optimised for each new waste stream or variation in composition encountered. Determining the optimal conditions for each new waste stream requires the collection of numerous experimental trials. This is resource intensive and presents a barrier to the deployment of yeast protein production processes.

Machine Learning (ML) methods offer an opportunity to improve the efficiency of fermentation optimisation. Data such as fermentation conditions (e.g., pH, temperature, inoculum density, time, aeration, and agitation), substrate composition (e.g., reducing sugars), and real-time sensor profiles (e.g., near-infrared spectra) have been utilised by ML models to predict product yield. These models have then been used to optimise process parameters and maximise the production of bioethanol, lactic acid, hydrogen, organic acids, and microbial enzymes (Said et al., 2023; Alloun and Calvio, 2024). ML models can be trained on experimental data to correlate substrate composition and process conditions to yeast growth. These models can then be used to predict yeast growth using new substrate compositions or under different conditions. This *in silico* method can guide targeted laboratory experimental trials to optimise the yeast production process for new waste streams with reduced time and resources. Specifically, hybrid modelling can be used to enhance the accuracy and generalisation capacity of these models by combining ML with traditional mathematical modelling.

Hybrid modelling has previously been used in fermentation processes to combine mechanistic mass balances, kinetic equations, and computational fluid dynamics simulations with empirical data or in-line sensing to predict substrate, biomass, and product concentrations over time. This can enable accurate real-time monitoring and control as well as the forecasting of fermentation behaviour at larger scale (Agharafeie et al., 2023; Albino et al., 2024). However, few studies have attempted to predict the productivity under different fermentation conditions and substrate compositions. For example, da Silva Pereira et al. (2021) developed a hybrid model for ethanol production from cashew apple juice by *S. cerevisiae*. Artificial neural networks (ANN) were used to predict the rates of substrate consumption, biomass growth, and product formation based on the inputs of current biomass and substrate concentration, temperature, and stirring speed. From this, it was determined that the ideal fermentation conditions should use an initial substrate concentration of 127 g/l, a temperature of 35 °C, an initial cell concentration of 5.8 g/l, and a stirring speed of 111 rpm. Pinto et al. (2022) used an ANN to predict mechanistic equation parameters based on inputs such as current substrate, biomass, and product concentrations as well as feed rates, temperature, and pH. The hybrid model was used to predict final biomass and product concentrations for a *Pichia pastoris* process expressing a single-chain antibody fragment. However, neither of these works attempted to optimise the fermentation process for new substrates or microbial strains.

To develop hybrid ML models able to optimise heterogeneous microbe strains and waste streams, a broad dataset is required opposed to relying on single, localised datasets as used in previous studies. In this work, aggregated datasets are used to enable generalised predictive accuracy over diverse waste streams. Experimental data was extracted from previously published literature utilising food waste streams to grow yeast biomass. Data from 963 fermentation experiments from 55 published articles spanning over 46 different yeast strains and 79 food side streams were used to train the hybrid ML model. The benefit of exploiting published experimental results is that the ML model can be trained on data across a diverse range of yeast strains, experimental procedures, food waste substrates, nutrient supplementation strategies, and fermentation conditions. The substrate compositions were extracted from the articles, estimated using referenced values, and included as model input features via textual embeddings.

The hybrid model correlated substrate composition and fermentation conditions to the final yeast biomass reported in the published articles. This was achieved by using the ML model to predict two Monod equation parameters (Monod, 1949), the maximum specific growth rate (μ*_max_*) and the half-saturation constant (*K_s_*). The Monod equation was used owing to its simplicity as an unstructured model and ability to describe growth across a wide range of microorganisms and conditions (Muloiwa et al., 2020). The utility of the hybrid model was demonstrated in a Bayesian Optimisation (BO) strategy where selected published articles were used as test sets. This proved the ability of the hybrid model to improve the optimisation efficiency of real, previously published datasets compared to traditional optimisation methods. The use of the hybrid model reduced the number of experiments required to optimise the yeast biomass production for these test sets. The hybrid model approach can therefore be used to simulate yeast biomass growth from new food side stream compositions or conditions and reduce the time and resources required to optimise yeast protein production processes. The novelty and contributions of this work are the compilation of a comprehensive dataset consisting of previously published experimental results (provided in the Supplementary Information as a downloadable Excel file), and the use of transfer learning by training the hybrid model on this diverse dataset to improve the efficiency of optimising yeast biomass growth on new strains and substrates.

## 2 Materials and methods

A flowchart outlining the integration of the Embedding model (Section 2.1.2), Hybrid model (Section 2.2), and BO (Section 2.3) is depicted in Figure 1. Textual data (e.g., yeast strain, food waste stream category) from the literature dataset is converted into a numerical representation through an Embedding model. This is combined with numerical data from the literature dataset (e.g., substrate composition, fermentation conditions) and used as input features in the hybrid model. To evaluate the effectiveness of the Hybrid model in enhancing the efficiency of yeast biomass production process optimisation, two BO strategies were evaluated: a conventional method without utilising the Hybrid model (Without Hybrid Model) and a method that uses the Hybrid model predictions to inform experimental trial selection (With Hybrid Model). The BO methods are compared using test sets selected from the published literature dataset and excluded from training of the Embedding model and initial training of the hybrid model.

**Figure 1:**
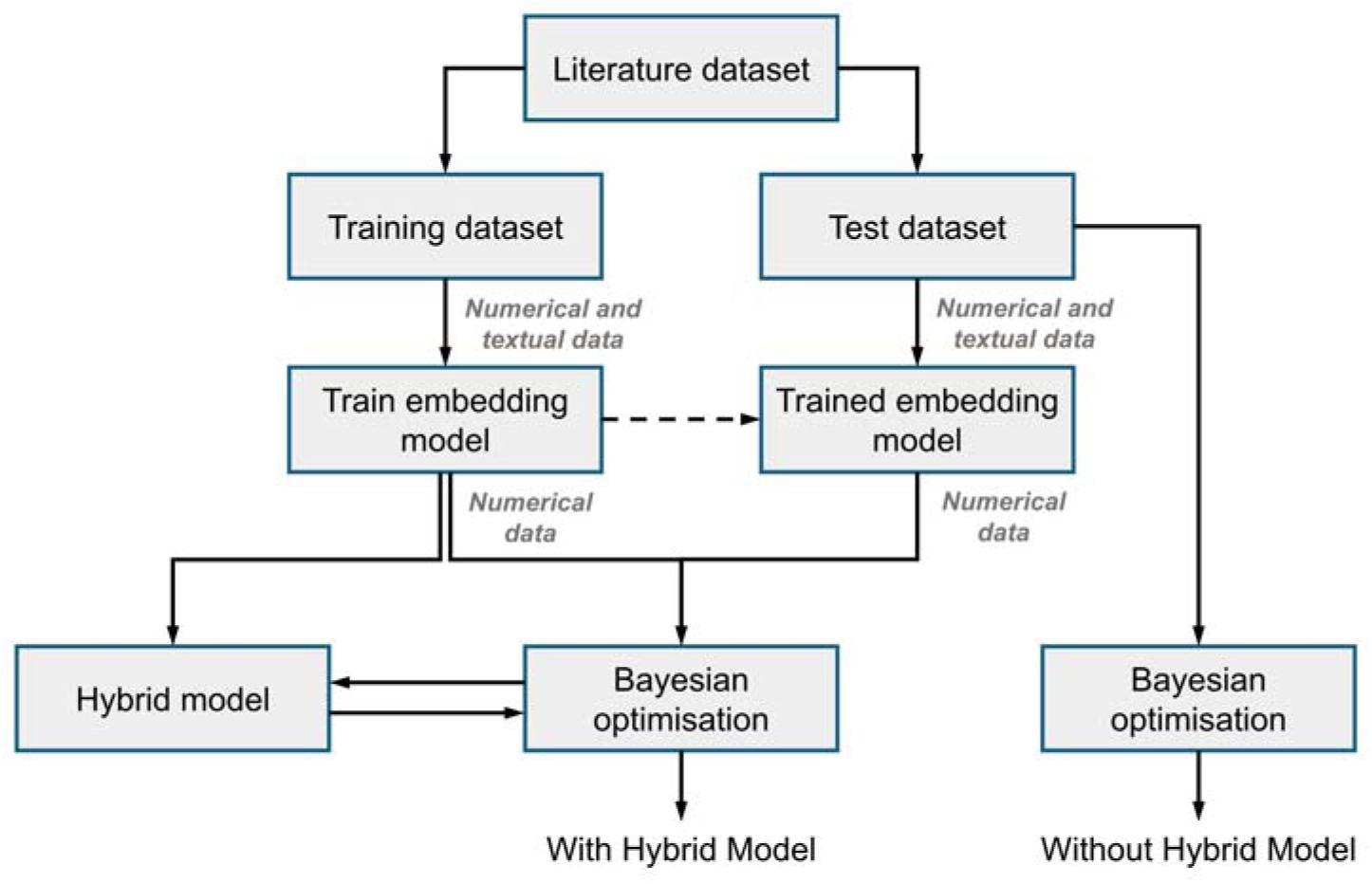
A flowchart outlining the integration of the Embedding model (Section 2.1.2), Hybrid model (Section 2.2), and BO (Section 2.3) within the methodology. The Embedding model is used to convert textual categories of yeast strains and waste streams into numerical representations for use as input features in the Hybrid model. Two BO methods were tested, a conventional method that did not utilise the Hybrid model predictions (Without Hybrid Model) and a method that did use the Hybrid model predictions to inform the selection of the next experimental trial (With Hybrid Model). The BO methods are evaluated on test sets selected from the published literature dataset and excluded from training of the Embedding model and initial training of the Hybrid model.

### 2.1 Data collection

The dataset was compiled using the existing published literature which utilised food waste streams as substrates for yeast biomass growth. A total of 963 fermentation experiments were included, derived from 55 publications. Overall, the dataset spans 46 yeast strains and 79 different food waste substrates. Compiling the data from diverse sources required several steps for integration of the dataset. Firstly, features across each study were collected for each published fermentation experiment (Table 1). Secondly, additional references were sought to estimate the substrate composition where this data was lacking from the published experimental fermentation data and are included in the downloadable Excel file dataset in the Supplementary Information. This allowed for inclusion of more fermentation experiments within the dataset and the creation of the Hybrid model able to correlate substrate composition to yeast biomass growth. Lastly, the varying supplementary nutrients used across the studies were converted to their compositional and elemental components (section 2.1.1 Supplementation input features).

**Table 1:**
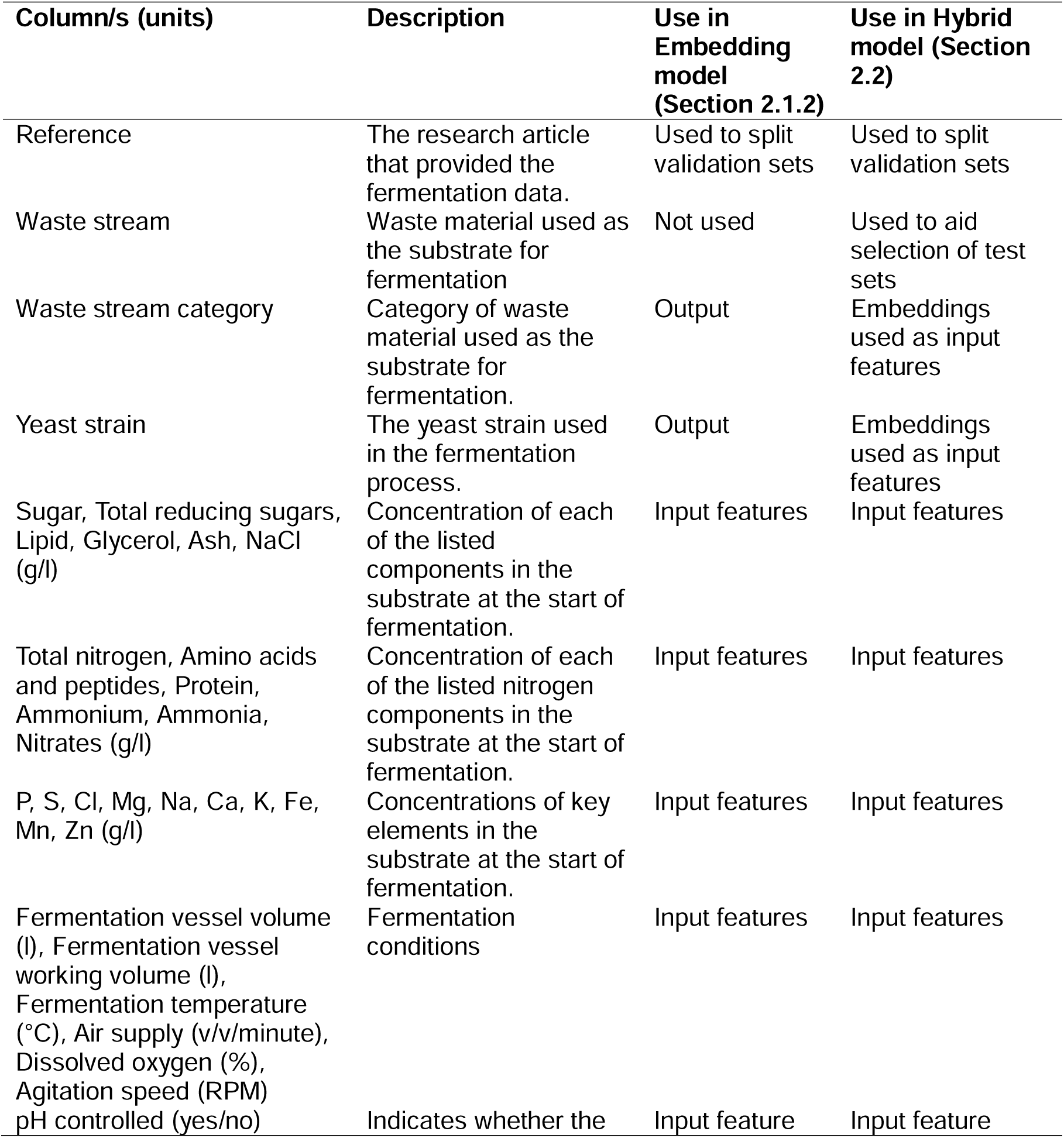
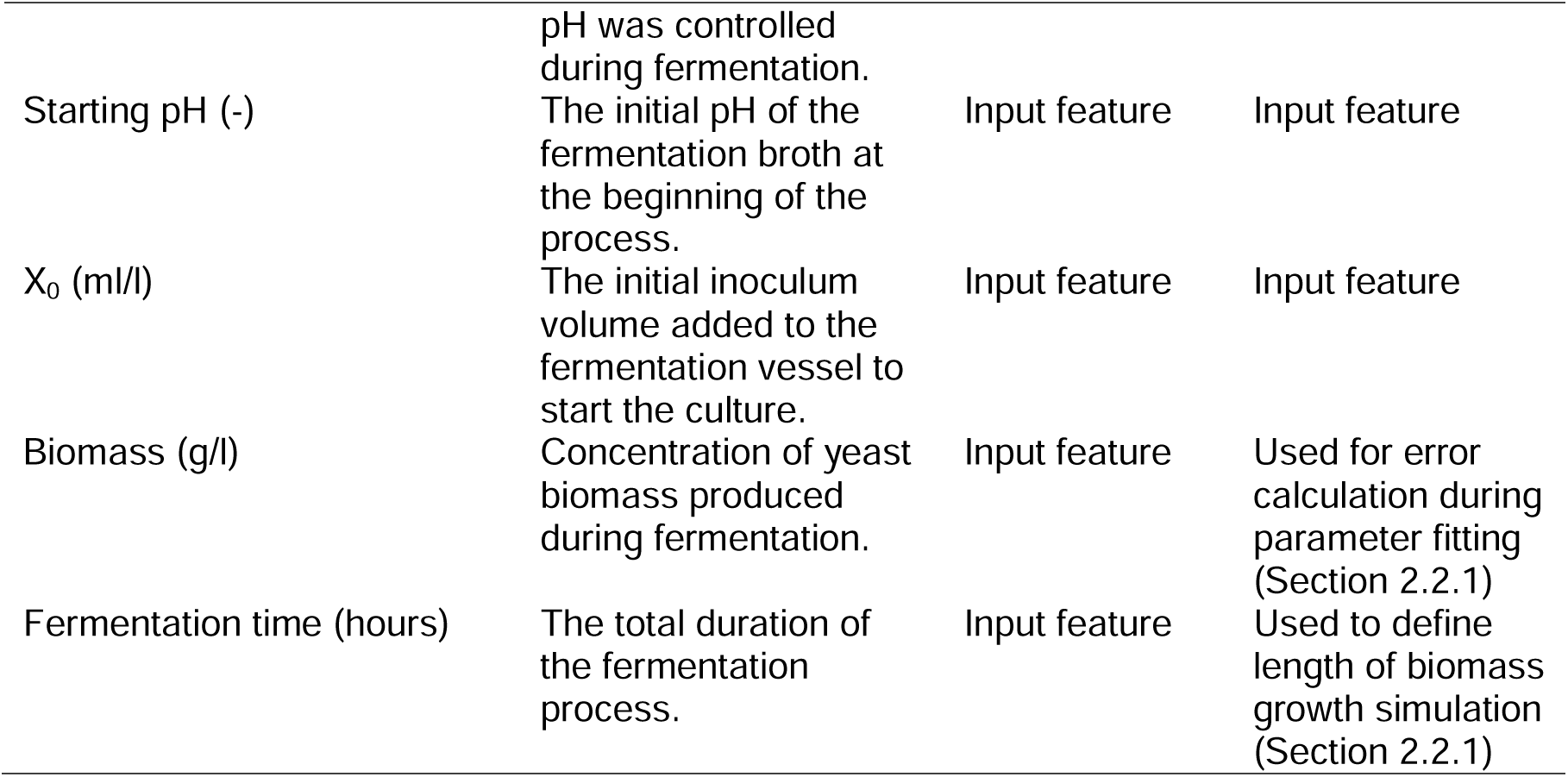
The data collected for each of the 963 fermentation experiments previously published in the literature.

#### 2.1.1 Supplementation input features

The supplements added to the fermentation substrates varied across the studies included in the dataset. To enable inclusion of nutrient supplements added during fermentation in the embedding and hybrid models, the various supplements were broken down into their elemental components (Table 1). For instance, magnesium sulphate was deconstructed into its sulphur and magnesium content, while urea was represented by its total nitrogen and ammonia components. For supplements without a universal composition, the reference data included in Table 2 was used. This allowed the supplement constituents to be used as continuous input feature variables opposed to categorical variables.

**Table 2:**
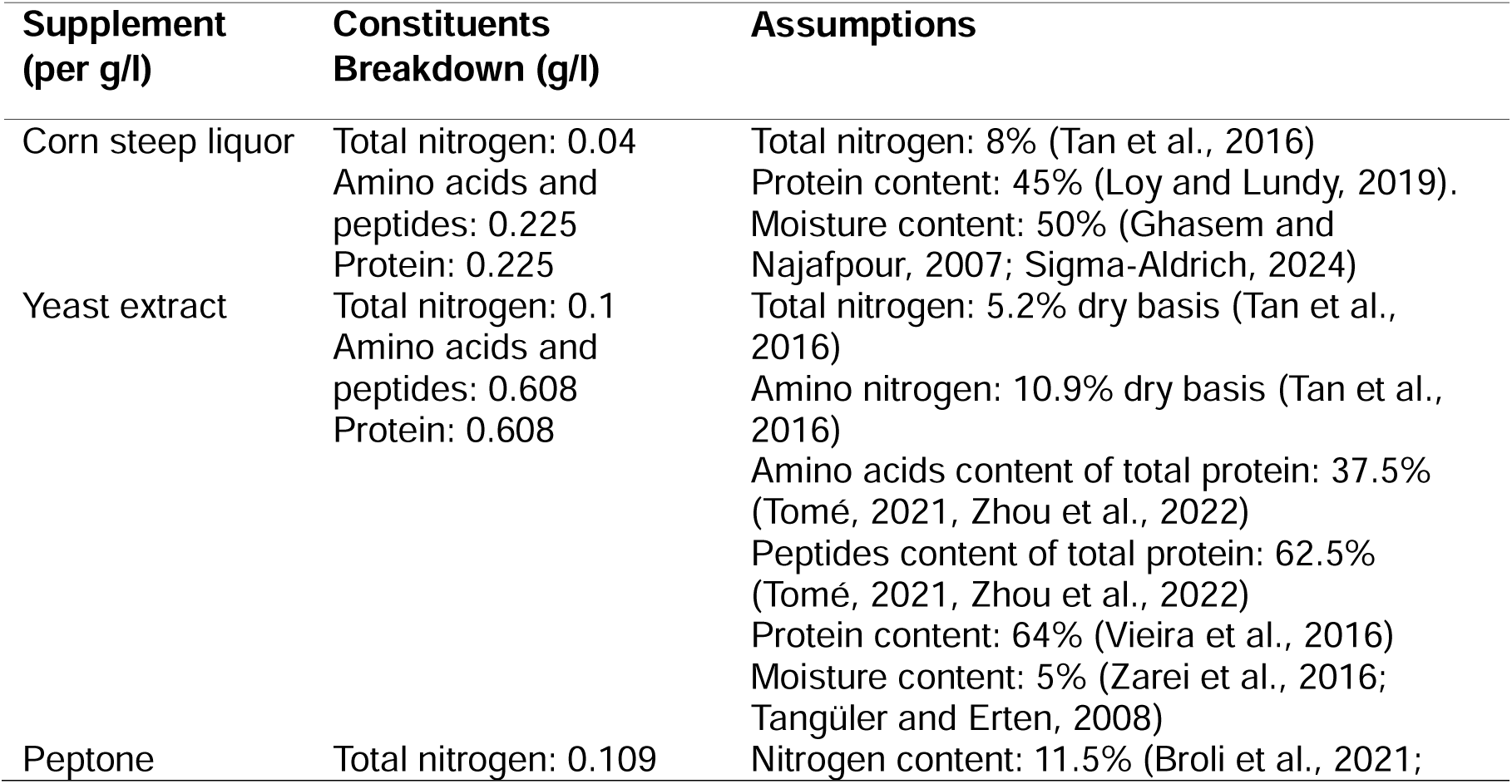
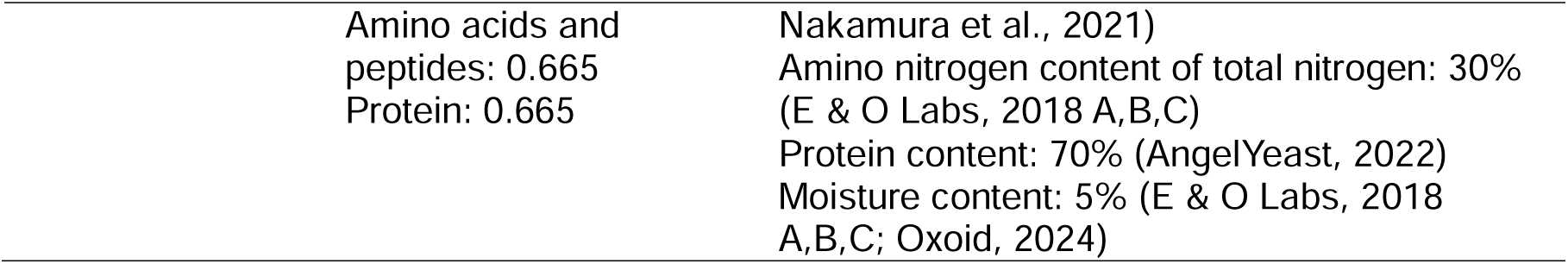
The constituent breakdown of supplements without a fixed composition. The breakdown enabled the supplements to be used as input features in the embedding and hybrid models (Table 1). Other supplements with fixed compositions (e.g., magnesium sulphate or urea) were also broken down into their constituents for use in the models.

#### 2.1.2 Embeddings for textual data

The waste stream category and yeast strain data columns (Table 1) were collected as textual data. An ANN-based Embedding model was used to convert these textual data columns into continuous numerical representations for use as input features in the hybrid model. This enabled the Hybrid model to learn relationships between the waste stream categories and yeast strains to better model their effects on the fermentation process. The Embedding model used all 33 input features from Table 1, excluding the textual columns (“Reference”, “Waste stream”, “Waste stream category”, and “Yeast strain”). The deep ANN used for the Embedding model was a multi-task model, trained to predict two outputs: the waste stream category and the yeast strain. This was a classification task using the cross-entropy loss where the Embedding model was trained to predict the correct categories for each fermentation data point. The model architecture consisted of two shared hidden layers (32 and 16 neurons), followed by two task-specific branches, each containing three hidden layers (8, 4, and 2 neurons). The ReLU activation function was used in all of the hidden layers. The first task-specific branch predicted the waste stream category while the second branch predicted the yeast strain.

The input features were standardised before training the Embedding model. Fermentation datapoints using mixed waste streams or yeast strains were excluded from the dataset (50 datapoints in total). For the BO trials (Section 2.3), the datapoints used in the test sets were also excluded from the dataset used to derive the embeddings. The Adam optimiser was used to train the model, with a learning rate of 0.001, for 20,000 epochs, using a batch size of 64. Five-fold cross-validation and grid search optimisation was employed to tune the ANN architecture and learning hyperparameters. The number of shared layers was trialled between one and three each with 2 to 128 neurons with a factor of two step size. Similarly, one to three task-specific layers were trialled consisting of 2 to 32 neurons with a factor of two step size. The learning rate was trialled between 10e^-1^ and 10e^-5^ with a factor of three step size, the batch size between 16 and 128 with a factor of two step size, and the number of epochs between 2,000 and 20,000 with a step size of 2,000. The validation sets were grouped based on their “Reference” column (Table 1) to avoid datapoints from the same published article being split between sets. The model was retrained on the full dataset after hyperparameter optimisation. After training, embeddings were extracted from the second-to-last fully connected layer (with two neurons), transforming the waste stream and yeast strain into numerical vectors that could be used in the Hybrid model (Figures 2 and 3).

**Figure 2:**
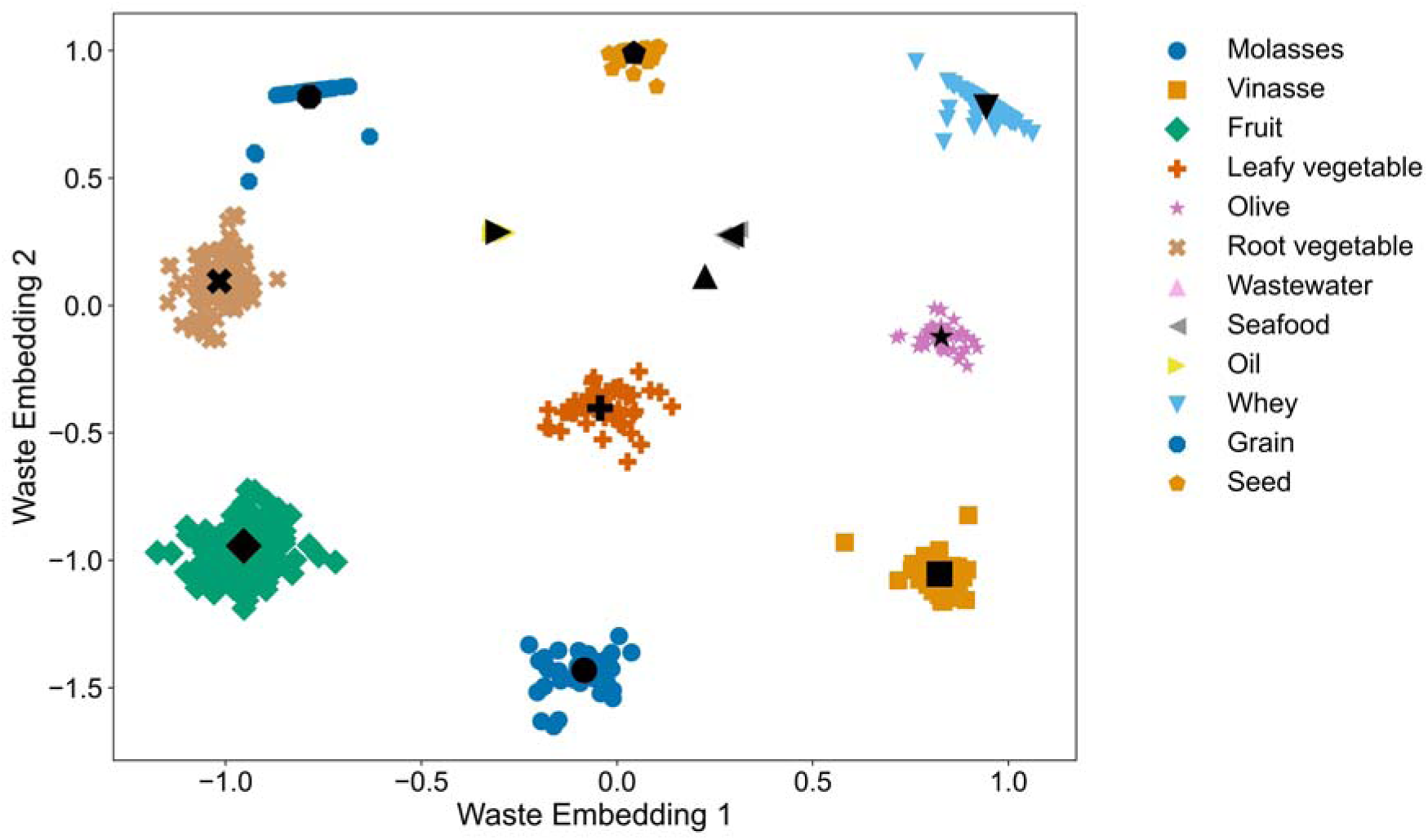
The numerical embeddings for each waste stream category learned by the Embedding model. The coloured markers indicate individual fermentation datapoints while the black markers indicate the centroid for all fermentation utilising that food waste stream. The centroid embeddings were used in the Hybrid model to numerically describe each waste stream.

**Figure 3:**
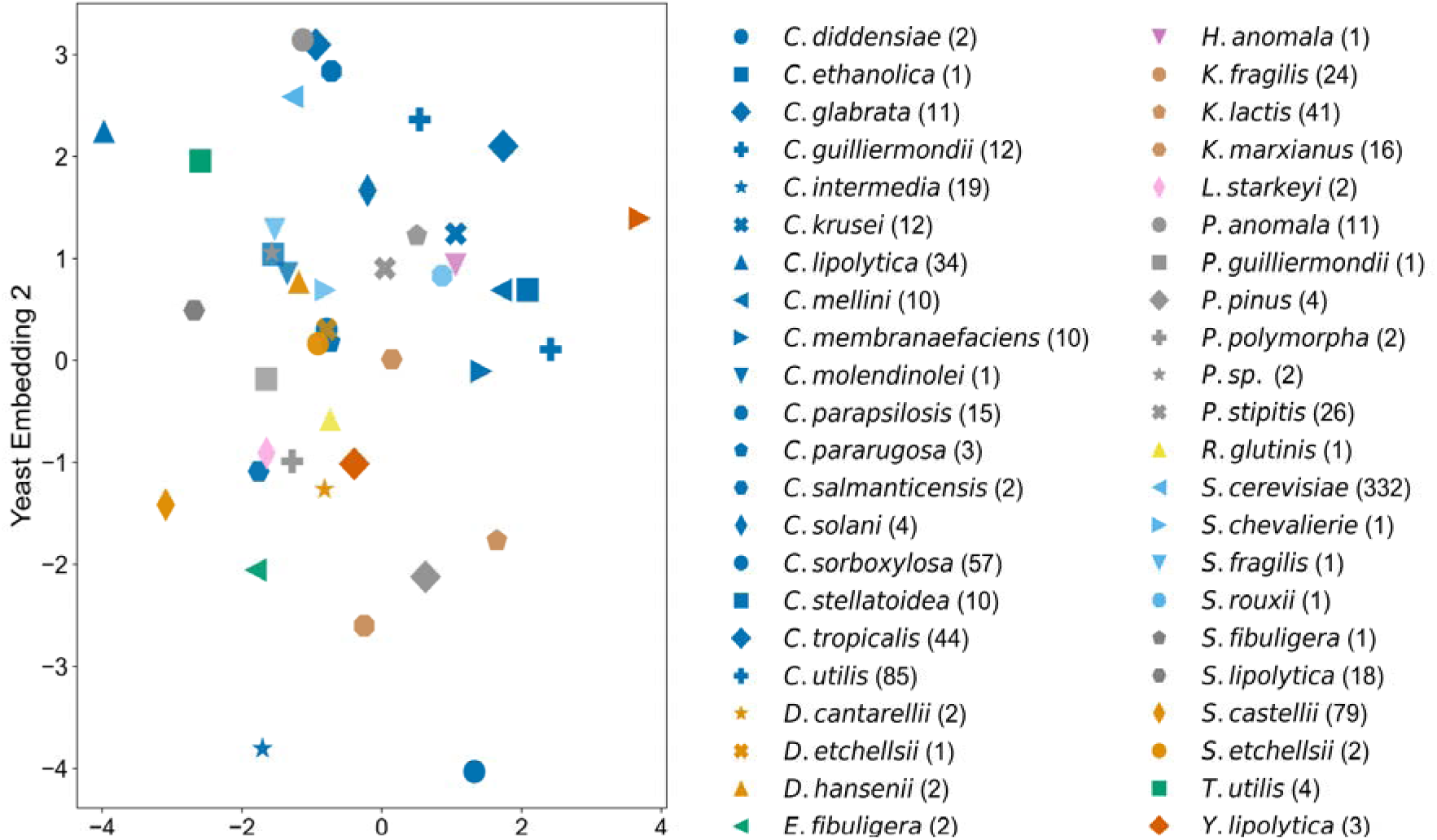
The numerical embeddings for each yeast strain learned by the Embedding model. The coloured markers indicate the centroid embeddings of the yeast strain. The individual fermentation datapoints were omitted to aid clarity. The centroid embeddings were used in the Hybrid model to numerically describe each yeast strain. The yeast genera include *Candida, Debaryomyces, Endomycopsis, Geotrichum, Hansenula, Kluyveromyces, Lipomyces, Pichia, Rhodotorula, Saccharomyces, Saccharomycopsis, Schwanniomyces, Torula,* and *Yarrowia*.

### 2.2 Hybrid modelling

Hybrid modelling combines traditional mechanistic models which rely on mathematical principles with data-driven models such as ML. The Hybrid model can then leverage the strengths of both approaches allowing the underlying dynamics of the mechanistic model to be matched to the real-world complexity via ML. In this work, Monod equation parameters were fit to the fermentation datapoints using the final yeast biomass concentrations extracted from the published articles (Section 2.2.1). Next, ML was used to correlate substrate composition and fermentation conditions to these Monod equation parameters (Section 2.2.2).

#### 2.2.1 Parameter optimisation

The Monod equation models the growth of microbial systems based on the concentration of a limiting substrate (for example, a type of sugar). Many extensions to the Monod equation have been proposed such as the inclusion of multiple substrates, inhibition terms, maintenance terms, Gaussian functions of temperature or pH, and additional lag and death stages (Bouguettoucha et al., 2011; Muloiwa et al., 2020). However, increasing the number of parameters requires increasing the dataset size to accurately fit each parameter. As the dataset is constructed from published literature, and therefore limited to the dataset size reported in each published article, the Monod equation with the fewest parameters was used to prevent over-fitting.

The Monod equation is expressed as:

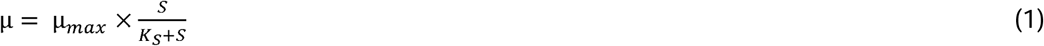

Where *µ* is the growth rate (h^-1^), *µ_max_*is the maximum specific growth rate (h^-1^), *K_S_* is the substrate half-saturation constant (g/l), and *S* is the substrate concentration (g/l). The growth biomass over time is described by:

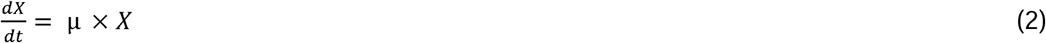

Where *X* (g/l) is the biomass concentration. Substrate depletion over time is described by:

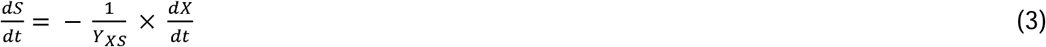

Where *Y_XS_*(g/g) is the yield coefficient, representing the amount of biomass produced per unit of substrate consumed.

In this study, the yield coefficient (Y_XS_) was fixed at 0.628, based on an analysis of the maximum biomass concentration reported in the collected dataset (62.48 g/l in Carsanba et al. (2023)) and the desired starting substrate range (0–100 g/l). A scaling factor, fixed at 0.001, was used to convert the starting biomass concentration, X_0_, (ml/l) taken from the data (Table 1) to a biomass concentration (g/l). The scaling factor was calculated by finding the smallest ratio of final biomass to starting X_0_ in the dataset (found to be approximately 0.001), representing no growth in biomass during this fermentation.

A grid search algorithm was implemented to fit the remaining parameters. The Monod equations were simulated over time using Python’s `odeint` function for the fermentation duration reported in the dataset (Fermentation time in Table 1). The summed squared error between the simulated and experimentally observed final biomass concentrations was minimised. The half-saturation constant, *K_s_*, was varied between 0 and 100 g/l with a step size of 1 g/l and constrained to be constant for all datapoints. The parameters *µ_max_*and *S_0_* (the starting substrate concentration) were optimized for each fermentation datapoint using a grid search approach. The search space for *µ_max_* ranged from 0 to 2 h^-1^ with a step size of 0.02 h^-1^, while the search space for *S_0_* ranged from 0 to 100 g/l with a step size of 1 g/l.

#### 2.2.2 Machine learning

The optimal Monod parameters identified for each experiment (Section 2.2.1) were used as target values for the ML model component of the hybrid model. The ML model was trained to predict the Monod equation parameters *µ_max_* and *S_0_* based on the input features derived from the fermentation dataset (Table 1). The model architecture consisted of a multi-task fully connected ANN with a single hidden layer consisting of 32 neurons. The ReLU activation function was used in the hidden layer. The output layer consisted of two neurons, corresponding to the predicted values of *µ_max_* and *S_0_*. The model was trained using the Adam optimiser, with a learning rate of 0.001 and a batch size of 32. Training was conducted for 10,000 epochs. The loss function used was the combined mean squared error between the predicted and observed values of *µ_max_* and *S_0_*. The input features were standardised before training. Five-fold cross-validation and grid search optimisation was employed to tune the ANN architecture and learning hyperparameters. The number of hidden layers was trialled between one and three each with 2 to 128 neurons with a factor of two step size. The learning rate was trialled between 10e^-1^ and 10e^-5^ with a factor of three step size, the batch size between 16 and 128 with a factor of two step size, and the number of epochs between 2,000 and 20,000 with a step size of 2,000. The validation sets were grouped based on their “Reference” column (Table 1) to avoid datapoints from the same published article being split between sets. The model was retrained on the full dataset after hyperparameter optimisation. The trained Hybrid model was used to simulate yeast growth for new substrate compositions and fermentation conditions during BO (Section 2.3). The datapoints used in each of the test sets were excluded from the datasets used to train the hybrid models.

### 2.3 Bayesian optimisation

The efficiency of optimising the substrate composition and fermentation conditions using the Hybrid model (With Hybrid Model) and when not using the Hybrid model (Without Hybrid Model) during BO was compared. Bayesian optimisation is a sequential optimisation strategy of functions that are expensive to evaluate such as experimental trials (Shahriari et al., 2015). The method uses a surrogate ML model with uncertainty estimation to learn the correlation between experimental parameters (e.g., substrate composition and fermentation conditions) and the objective function (e.g., maximising yeast biomass growth). The objective function is the output that an optimisation process is aiming to maximise or minimise. An acquisition function is employed to select the next experimental trial to conduct by balancing exploration, selection of parameters with high uncertainty, and exploitation, selection of parameters with a high predicted objective function value.

To evaluate the utility of the hybrid modelling method, five published articles from the dataset were selected as test sets and excluded from the training of the embedding and hybrid models. The articles were selected based on their recency, large number of fermentation experiments conducted, and diversity of yeast strain and food waste stream trialled (Table 3). The five selected studies date from 2019 to 2023 (the most recent time period included within the full dataset which spans from 1975 to 2023) to reflect the latest research within the area. The selected studies also contain the largest (Dos Reis et al., 2019), third largest (Sawsan et al., 2021), and fifth largest (Diboune et al., 2019) individual datasets within the full collated dataset. This enables the hybrid modelling method to be evaluated on the largest optimisation problems available. These studies also included new waste substrates and yeast strains not included in the training dataset. For example, Sawsan et al. (2021) used grape juice, Rages et al. (2021) used olive waste, and Diboune et al. (2019) used opuntia ficus indica pulp to grow yeast biomass. Furthermore, Dos Reis et al. (2019) trialled *Candida glabrata* and *Pichia anomala* to ferment vinasse. Inclusion of these datasets enabled evaluation of whether the hybrid model could efficiently optimise process parameters outside of its training dataset. To achieve this, grape juice and opuntia ficus indica pulp were assigned the waste embeddings of the fruit category and olive waste was assigned the embeddings of the olive category (Figure 2). To derive the yeast embeddings for the new yeast strains (Figure 3), the datapoints used to initialise the BO method for Dos Reis et al. (2019) were passed through the embedding model.

**Table 3:**
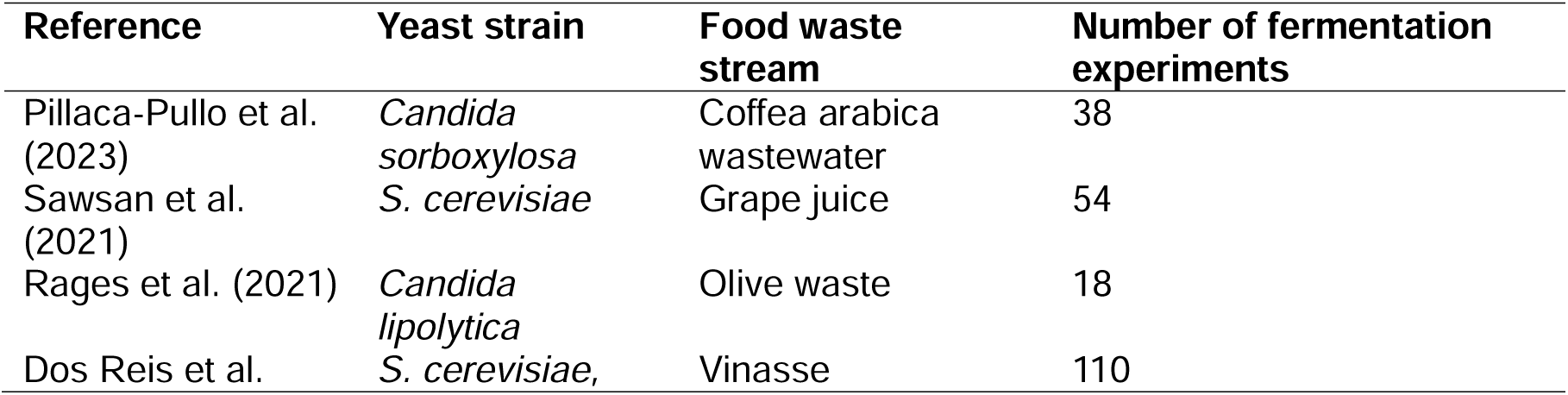
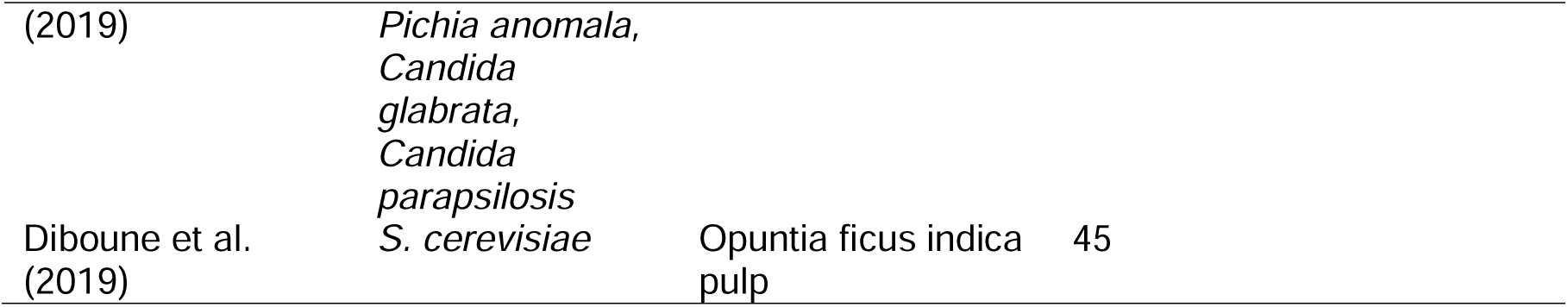
The five published articles used as test sets to evaluate the utility of the Hybrid model. The articles were selected based on their recency, large number of fermentation experiments conducted, and diversity of yeast strain and food waste stream trialled. These articles were excluded from the training of the Embedding model and the initial Hybrid model. The efficiency required to optimise the substrate composition and fermentation conditions of these test sets was compared both with (With Hybrid Model) and without using the Hybrid model (Without Hybrid Model) during BO.

During BO, both the With Hybrid Model and Without Hybrid Model methods were constrained to select trials from the combinations of parameters (e.g., yeast strain, supplementation, conditions) trialled within the test set articles. Both methods were initialised using the three parameter combinations trialled in the test set articles that produced the smallest final biomass concentrations. This represents the most difficult optimisation task possible for the two methods given the lowest possible number of starting datapoints and objective function values. The Without Hybrid Model method used only the results of the trialled experiments to select the next parameter combination to trial via the acquisition function. The With Hybrid Model method used the trialled experiment results in conjunction with the final yeast biomass concentrations predicted by the trained Hybrid model predictions for the remaining untrialled parameter combinations. Once a new parameter combination was trialled, the Hybrid model was retrained to include this datapoint and used to aid the next parameter combination selection.

The Random Forest (RF) algorithm was used as the surrogate model during BO to correlate the substrate composition and fermentation conditions to the final yeast biomass concentration. Random Forest is an ensemble algorithm and thereby enables the prediction variance to be used as the exploration component of the acquisition function. K-fold cross-validation, where K begins as three during initialisation and increases to five as further experiments were trialled, and grid-search optimisation was used to tune the RF hyperparameters. The Expected Improvement acquisition function was used owing to it being the most commonly used in BO (Shahriari et al., 2015).

## 3 Results and discussion

The performances of the With Hybrid Model and Without Hybrid Model methods during BO of the five test sets are displayed in Figure 4. The use of the test sets proved the ability of the With Hybrid Model to improve the efficiency of fermentation optimisation on real, previously published datasets. The With Hybrid Model methodology exhibited enhanced efficiency in optimising all test sets compared to the Without Hybrid Model strategy. This shows that hybrid ML-BO framework (With Hybrid Model) outperforms conventional optimisation relying on laboratory data collection alone (Without Hybrid Model). By transferring the knowledge learned from the literature-derived dataset, the With Hybrid Model approach decreases the number of laboratory experiments needed to determine optimal conditions for yeast biomass production from food waste substrates.

**Figure 4:**
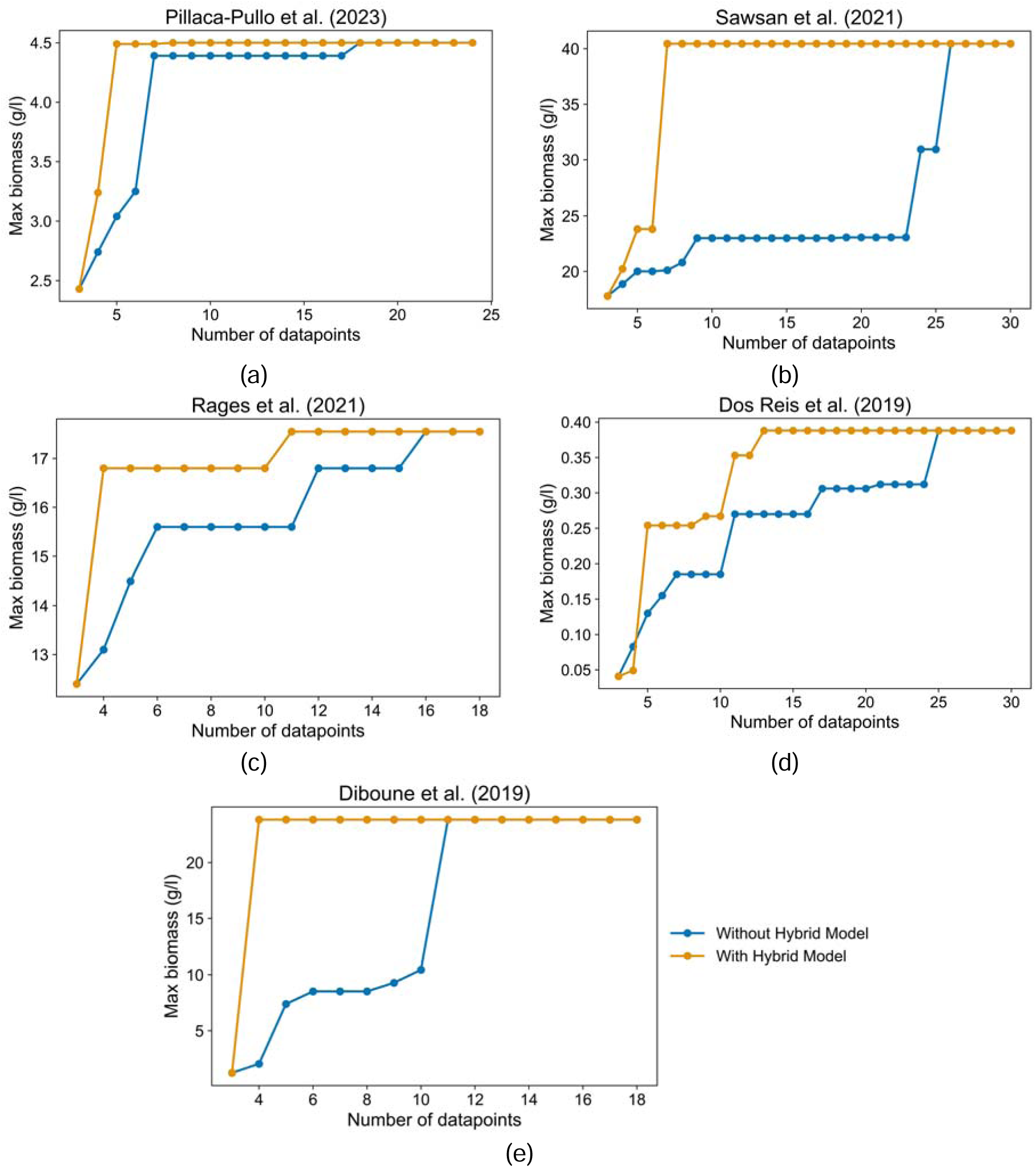
The performance of the Without Hybrid Model methods (blue) and With Hybrid Model (orange) during BO of the five test sets. The With Hybrid Model methodology required fewer datapoints (experimental trials) to optimise the fermentation processes for all test sets evaluated compared to the Without Hybrid Model strategy. (a) Pillaca-Pullo et al. (2023), (b) Sawsan et al. (2021), (c) Rages et al. (2021), (d) Dos Reis et al. (2019), and (e) Diboune et al. (2019).

On average, the With Hybrid Model required 10.6 fewer data points (experimental trials) to optimise the fermentation systems, with reductions ranging from 5 fewer data points in Rages et al. (2021) to 19 fewer in Sawsan et al. (2021). For Rages et al. (2021), the BO was initialised with three experimental trials containing 2 g/l peptone. Although the optimal substrate composition utilised 4 g/l peptone, higher concentrations demonstrated inhibitory effects. When the With Hybrid Model method selected trials with 5 to 7 g/l peptone, it resulted in lower yeast biomass concentrations than the optimal conditions. This necessitated additional data collection for the With Hybrid Model, yet it still required fewer data points than the Without Hybrid Model method. Conversely, in Sawsan et al. (2021), one of the initial experimental trials matched the optimal conditions in sugar, urea, ammonium sulphate, and inoculum concentrations as well as temperature but used a more acidic starting pH (0.951 compared to the optimal 4.5). This condition severely hindered the Without Hybrid Model method, as the BO could not determine which variables contributed to the reduced yeast biomass concentration without extensive data collection. The With Hybrid Model method, leveraging its prior knowledge from the published literature, was able to identify the optimal conditions much more quickly.

On average, the With Hybrid Model required 5.6 additional data points after initialisation, ranging from one additional data point for Diboune et al. (2019) to ten for Dos Reis et al. (2019). In contrast, the Without Hybrid Model method required an average of 16.2 additional data points, ranging from eight for Diboune et al. (2019) to 23 for Sawsan et al. (2021). In the case of Diboune et al. (2019), the With Hybrid Model optimised the fermentation system with only one additional data point because the optimal substrate composition uniquely included yeast extract. The Without Hybrid Model method was unable to learn the relationship between yeast extract addition and yeast biomass growth due to its absence in all other experimental trials, whereas the With Hybrid Model utilised its knowledge from existing literature to predict the impact of including yeast extract. In Dos Reis et al. (2019), a greater number of additional data points were required, attributed to the dataset’s size (110 data points) and the inclusion of four yeast strains (*S. cerevisiae*, *P. anomala*, *C. glabrata*, and *C. parapsilosis*). Overall, the With Hybrid Model method achieved a 66% reduction in the number of experimental trials needed compared to the Without Hybrid Model method, with reductions varying from 39% in Rages et al. (2021) to 88% in Diboune et al. (2019).

To interpret the relationships learned by the Hybrid model between the substrate composition and fermentation conditions on the yeast growth parameters, the starting substrate concentration parameter (*S_0_*) and the maximum specific growth rate parameter (*µ_max_*), SHAP values (SHapley Additive exPlanations) were used (Figures 5a and 5b). This is a ML explainability method that quantifies the contribution of each input feature to the outputs. SHAP values consider all possible combinations of feature contributions to calculate an average marginal effect for each input (Molnar, 2022). SHAP values reveal not only the magnitude but also the direction (positive or negative) of the impact of each input feature on the yeast growth parameters.

**Figure 5:**
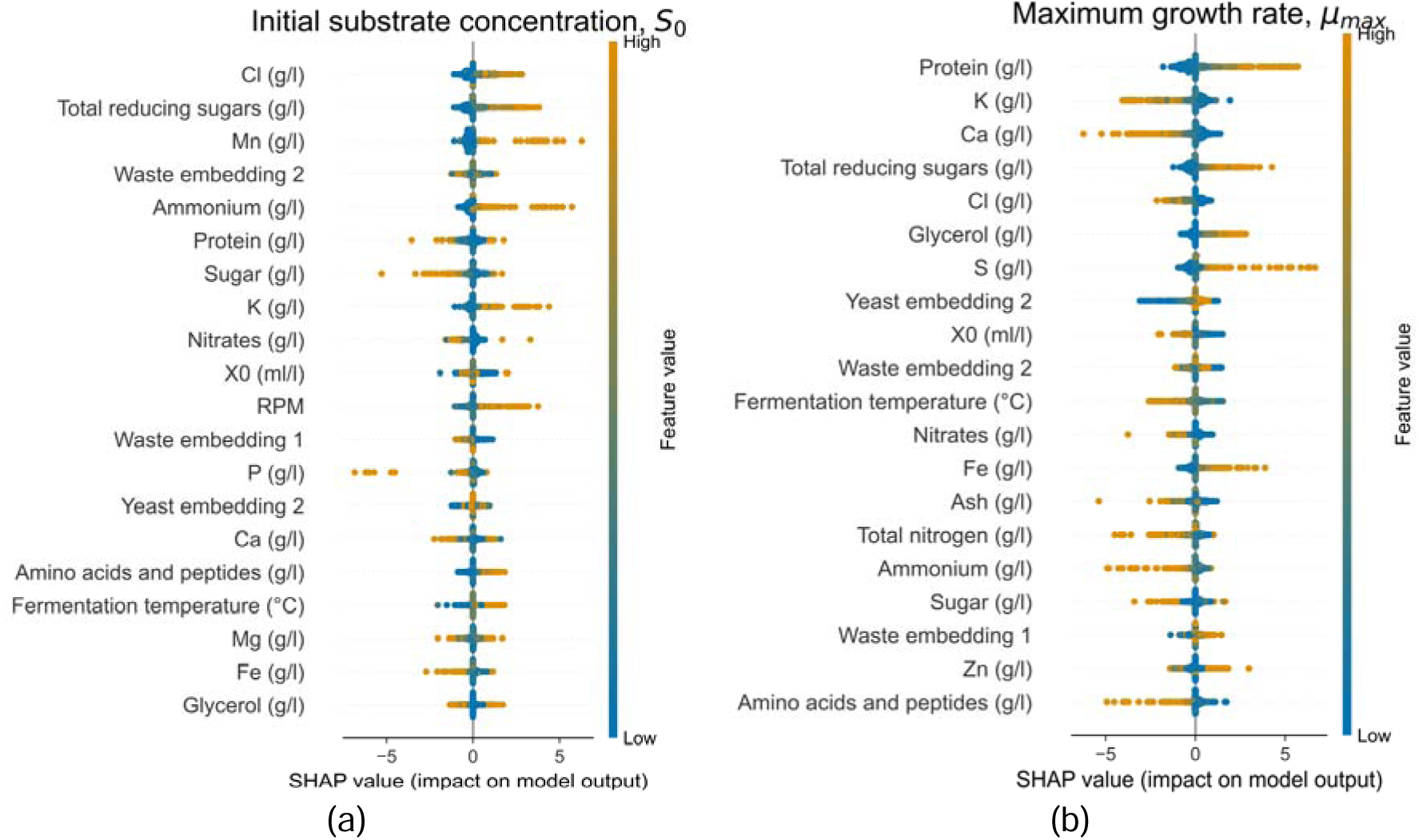
The SHAP plots illustrating the feature importance for the Hybrid model to predict yeast growth parameters: (a) The initial substrate concentration, *S_0_*. (b) The maximum specific growth rate, *µ_max_*.

Chloride, manganese, and potassium concentrations were all shown to positively affect the initial substrate concentration parameter (*S_0_*). However, while potassium and chloride enhance S, they were found to negatively impact the maximum specific growth rate parameter (*µ_max_*). This is likely due to the inhibitory influence of high chloride ion concentrations on microbial growth rates, despite their role in enhancing enzyme activity and facilitating membrane transport for improved substrate uptake (Belz et al., 2017; Jennings and Cui, 2008). Similarly, potassium is essential for maintaining osmotic balance and enzyme activation but becomes inhibitory to yeast growth at elevated concentrations (Kahm et al., 2012; Barreto et al., 2012). Manganese, acting as a cofactor for numerous enzymes and enhancing enzyme activity to improve substrate utilisation efficiency (Thines et al., 2019), did not reach inhibitory levels in the studied literature dataset, maintaining its positive effect on *S_0_* without negatively affecting *µ_max_*. Conversely, calcium concentration was found to negatively impact *µ_max_*, suggesting it was mainly inhibitory to yeast growth within the published literature dataset.

Higher concentrations of reducing sugars were associated with increases in both *S_0_* and *µ_max_*owing to their role as readily metabolizable carbon sources for many yeast strains. This also indicates that reducing sugars were not present at concentrations high enough to inhibit yeast growth in the dataset. In contrast, concentrations of ammonium, amino acids, and peptides positively affected *S_0_* but negatively impacted *µ_max_*. Despite being readily assimilable nitrogen sources essential for protein synthesis and microbial growth, these components became inhibitory to yeast growth at high concentrations. Nitrate concentration showed negative impacts on both *S_0_* and *µ_max_*, indicating it is a less efficient nitrogen source compared to ammonium, amino acids, and peptides, likely due to the energy-intensive reduction steps required for nitrate assimilation into amino acids (Sanz-Luque et al., 2015). Protein concentration exhibited no clear impact on *S_0_* but had a positive effect on *µ_max_* suggesting that protein levels in the literature dataset were not high enough to inhibit yeast growth.

Revolutions per minute (RPM) displayed a slight positive impact on *S_0_*, likely because increased agitation rates improve oxygen transfer and homogenisation of the culture medium, facilitating substrate accessibility. Glycerol, sulphur, and iron concentrations were all shown to positively affect *µ_max_*, indicating that their levels were below the threshold that would inhibit yeast growth. Glycerol can serve as an additional carbon source and plays a role in osmoregulation (Blomberg, 2022); sulphur is a component of amino acids like cysteine and methionine, supporting protein synthesis (Thomas and Surdin-Kerjan, 1997); and iron is critical for electron transport and energy metabolism (Lindahl et al., 2020; Ramos-Alonso et al., 2020). The Yeast Centroid 2 embedding value was found to positively impact *µ_max_*. Well-studied yeast strains such as *S. cerevisiae* and *C. utilis* produced high Yeast Centroid 2 embedding values (Figure 3). This is likely due to these commonly used strains having higher growth rates and optimal growth conditions that are easier to reproduce, given their prevalence in scientific literature. Overall, the results from the SHAP analysis indicated that reducing sugar, manganese, protein, glycerol, sulphur, and iron concentrations as well as agitation can all be used at above average rates compared to the literature dataset to enhance yeast growth. In contrast, calcium and nitrate concentrations should be reduced compared to the levels used in the published literature owing to their inhibitory effects on yeast growth.

## 4 Future work

In this work, the Monod equation was used as it is an unstructured kinetic model that relates microbial growth rate to the concentration of a single limiting substrate without detailing internal metabolism (Du et al., 2022). However, the presence of the ML component within the Hybrid model changes the modelling strategy. Since ML can learn patterns directly from data, the mechanistic model need not be exhaustive. For example, the concentration of the limiting substrate, *S_0_*, was predicted using ML based on the substrate composition, yeast strain, and fermentation conditions making it a pseudo-substrate parameter (Table 1). This means that uncertain aspects (complex substrate composition, minor inhibitory effects) could be learned by the ML component. The Hybrid model therefore works with the mechanistic part enforcing basic scientific constraints to yield physically realistic outputs and the ML component fits to the remaining uncertainty. Furthermore, Monod kinetics have been widely used as a basis for modelling fermentation and have few parameters making it easy to fit with limited data (González-Figueredo et al., 2018). Some of the datasets collected from previous literature only contained a single datapoint and so the Monod equation, being the simplest model that incorporates core microbial dynamics, was utilised to prevent over-fitting to unnecessary parameters within the model.

However, more complex unstructured models can build on Monod kinetics by incorporating additional terms for phenomena that can affect growth. For example, at very high substrate concentrations, growth can slow down due to inhibitory effects. Models like the Andrews equation add an inhibition term to Monod kinetics to incorporate the decrease in the growth rate (Du et al., 2022). Similarly, product inhibition models can include terms for accumulated by-products. Microbial growth rate is also strongly influenced by temperature, pH, and dissolved oxygen, often following an optimum curve where if these parameters are too low or too high growth is impaired. Additional terms could be added that multiply the growth rate by a Gaussian function to represent parameter dependence (Jin et al., 2022). First order death rate equations can also be added to simulate biomass decline. Another way to model microbial growth is via metabolic models and Flux Balance Analysis (FBA), which are structured approaches (Du et al., 2022). Instead of describing growth with an empirical rate law, FBA uses a metabolic network for the microbe and calculates how nutrients are converted to biomass and products. However, to build a FBA model, detailed metabolic pathways of the yeast strain are needed. While these models exist for common strains such as *S. cerevisiae* on ideal media, applying FBA to less common strains and using food waste substrates introduces complexity as the waste is a mixture of many components (Table 1) and can reduce the prediction accuracy of growth rates. On the other hand, the most important pathways could be selected (e.g., carbon) and ML can be used to fit the model to experimental data in a hybrid modelling approach. Future work can compare the use of these more complex models with greater numbers of parameters. To prevent overfitting the selection of model complexity can form part of the hybrid model validation procedure, selecting the mechanistic models that maximise accuracy on the validation set after training.

Future work will also focus on strengthening the Hybrid model’s ability to accurately predict microbial growth parameters for larger scale fermentations. While the dataset already spans working volumes from 0.05 to 10,000 litres, the number of data points at this higher capacity were limited compared to small– and bench-scale fermentations (Figure 6). Additional fermentation data from large-scale processes that did not use food wastes substrates could be added to the dataset. This would enable the Hybrid model to learn how parameters likely to change across scales (e.g., agitation speed and method) affects yeast growth. Furthermore, collection of data at pilot-scale can be added to the dataset and the model retrained. This would fit the Hybrid model to this larger scale data while suggesting experimental trials within a Bayesian optimisation approach. The Hybrid model could also be fine-tuned for different scales, such as giving larger loss weightings to datapoints at the desired experimental scale.

**Figure 6:**
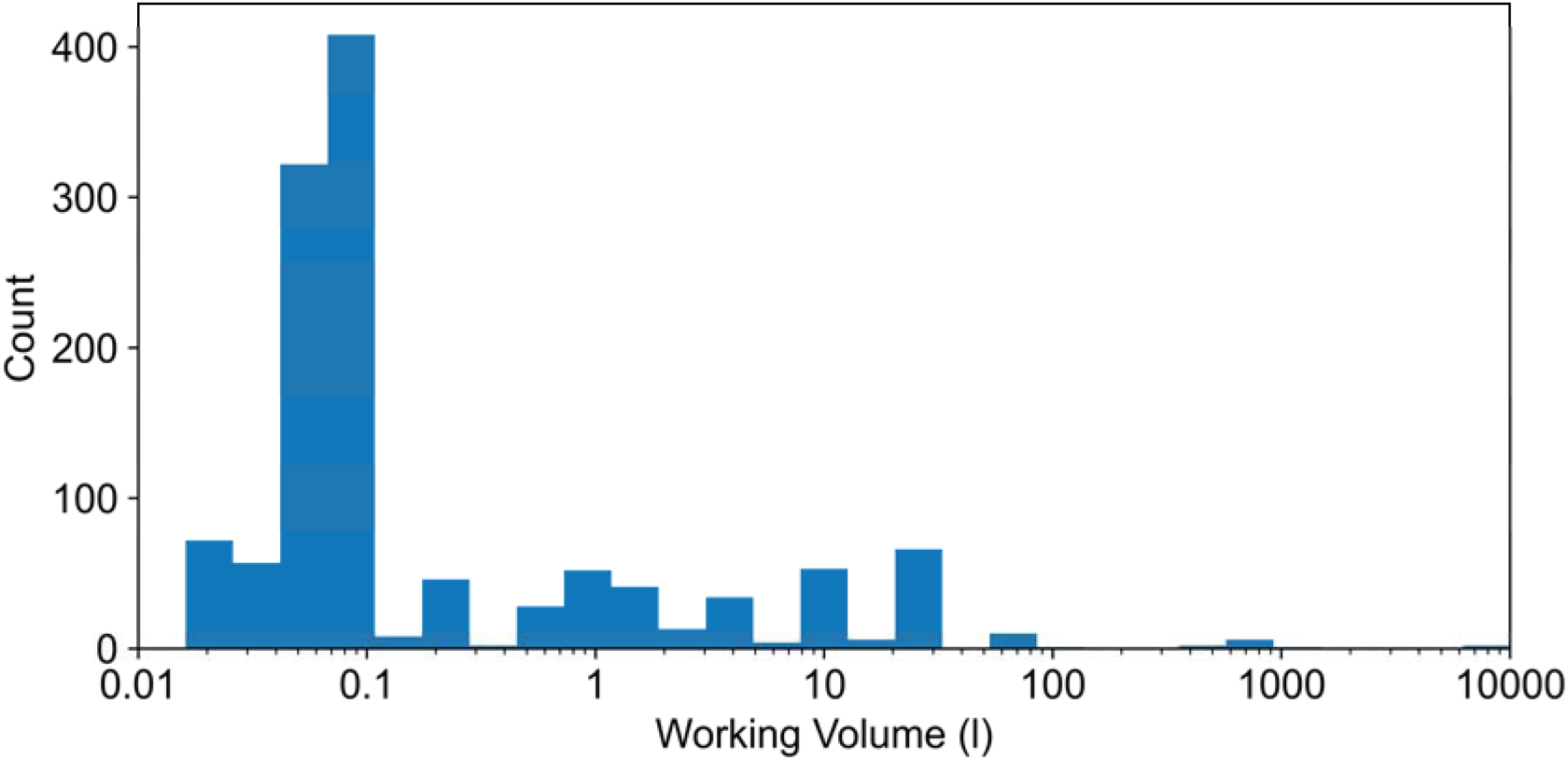
The distribution of fermentation vessel working volumes included in the dataset (available as a downloadable Excel file in the Supplementary Information).

## 5 Conclusions

In this study, a hybrid ML approach was employed to enhance the efficiency of fermentation process optimisation using BO for yeast biomass growth using food waste streams as substrates. The utility of the Hybrid model was demonstrated in a BO strategy where selected published articles were used as test sets. This proved the ability of the Hybrid model to improve the optimisation efficiency of real, previously published datasets compared to traditional optimisation methods. The utilisation of the Hybrid model led to an average reduction of 66% in the number of experimental trials required to identify the global maximum of yeast biomass during BO compared to conventional methods. Across the test sets evaluated, reductions ranged from 39% to 88%. On average, when optimising using the Hybrid model 5.6 additional experimental trials were required after initialisation, compared to 16.2 additional fermentation trials without the Hybrid model approach. The novelty and contributions of this work are the collection and utilisation of an extensive dataset comprising previously published experimental results, which is provided in the Supplementary Information as a downloadable Excel file; and the use of transfer learning by training the Hybrid model on this dataset to improve the efficiency of optimising yeast biomass growth on new strains and substrates.

The Hybrid model was developed using a dataset comprising 963 fermentation experiments sourced from 55 publications, encompassing 46 yeast strains and 79 food waste substrates. The model integrates an ML component that correlates yeast strains, substrate compositions, and fermentation conditions with two Monod growth parameters: the initial substrate concentration (*S*) and the maximum specific growth rate (*µ_max_*). An Embedding model was used to convert categorical data related to waste streams and yeast strains into numerical input features for the ML model. Yeast growth dynamics were simulated over time using the Monod equations, enabling the prediction of final biomass concentrations. By combining mechanistic models with data-driven ML techniques, the Hybrid model leverages the strengths of both approaches, aligning theoretical dynamics with the complexities of real-world data. Analysis using SHAP indicated that higher concentrations of reducing sugar, manganese, protein, glycerol, sulphur, and iron, as well as increased agitation, could be used than levels present in the published literature dataset to enhance yeast growth. Conversely, lower concentrations of calcium and nitrate than in the previous literature could be used due to their inhibitory effects on yeast biomass production.

## CRediT authorship contribution statement

**Alexander L. Bowler**: Conceptualization; Data curation; Formal analysis; Funding acquisition; Investigation; Methodology; Software; Validation; Visualization; Roles/Writing – original draft; and Writing – review & editing. **Nasser Alkhulaifi**: Software; Visualization; Roles/Writing – original draft; and Writing – review & editing. **Sarah Rodgers**: Data curation; Methodology; Software; Visualization; Roles/Writing – original draft; and Writing – review & editing. **Joanna H. Sier**: Conceptualization; Funding acquisition; Methodology; Supervision; Writing – review & editing. **Célia Ferreira**: Conceptualization; Funding acquisition; Methodology; Supervision; Writing – review & editing. **Darren Greetham**: Conceptualization; Funding acquisition; Methodology; Supervision; Writing – review & editing. **Jordan Pennells**: Conceptualization; Funding acquisition; Methodology; Writing – review & editing. **Kai Knoerzer**: Conceptualization; Funding acquisition; Methodology; Writing – review & editing. **Nicholas J. Watson**: Funding acquisition; Project administration; Resources; Supervision; Writing – review & editing.

## Acknowledgements

This work was supported by the BBSRC-funded project “AI-Optimised Fermentation for Sustainable Protein Production from Food Side Streams” [Grant number: BB/Y513933/1].

## Appendix A. Supplementary material

### Declaration of Generative AI and AI-assisted technologies in the writing process

During the preparation of this work the authors used OpenAI o1-preview to improve the writing’s readability once an advanced draft text had been established. After using this tool/service, the authors reviewed and edited the content as needed and takes full responsibility for the content of the publication.

